# Evolution of β-lactamase mediated cefiderocol resistance

**DOI:** 10.1101/2022.01.29.478156

**Authors:** Christopher Fröhlich, Vidar Sørum, Nobuhiko Tokuriki, Pål Jarle Johnsen, Ørjan Samuelsen

## Abstract

Cefiderocol is a novel siderophore β-lactam with improved hydrolytic stability toward β-lactamases, including carbapenemases, achieved by combining structural moieties of two clinically efficient cephalosporins, ceftazidime and cefepime. Consequently, cefiderocol represents a treatment alternative for infections caused by multi-drug resistant Gram-negatives. Using directed evolution on a wide variety of different β-lactamases, such as KPC-2 and CTX-M-15 (Ambler class A), NDM-1 (class B), CMY-2 (class C) and OXA-48 (class D), we studied the role of cefiderocol during β-lactamase-mediated resistance development. First, we investigated how the expression of different β-lactamases causes changes in cefiderocol susceptibility. In a low-copy number vector, we found that OXA-48 and KPC-2 conferred non or marginal decreases in cefiderocol susceptibility, respectively. On the contrary, CMY-2, CTX-M-15 and NDM-1 substantially decreased cefiderocol susceptibility by 16-, 8- and 32-fold, respectively. Second, we determined the evolutionary potential of these enzymes to adapt to increasing concentrations of cefiderocol. Our data show that with the acquisition of only 1 to 2 mutations, all β-lactamases were evolvable to further cefiderocol resistance by 2- (NDM-1, CTX-M-15), 4- (CMY-2), 8- (OXA-48) and 16-fold (KPC-2). Cefiderocol resistance development was often associated with collateral susceptibility changes including increased resistance to ceftazidime and ceftazidime-avibactam as well as functional trade-offs against different β-lactam drugs. Taken together, contemporary β-lactamases of all Ambler classes can potentially contribute to cefiderocol resistance development and can acquire mutations allowing them to adapt to increasing cefiderocol concentration. At the same time, resistance development caused clinically important cross-resistance, especially against ceftazidime combinations.

**Summary:** Despite the reported higher stability of cefiderocol against β-lactamase hydrolysis, we show that the expression of β-lactamases from different Ambler classes significantly contributes to cefiderocol resistance and that these enzymes have the evolutionary potential to evolve towards increasing cefiderocol concentrations.

## INTRODUCTION

The novel and recently introduced cephalosporin cefiderocol is a promising treatment option for infections caused by multi-drug resistant and carbapenemase-producing Gram-negatives based on two distinctive structural features [1]. Firstly, the cephalosporin molecule is linked to a catechol moiety (siderophore) promoting binding of iron, and thus facilitating uptake through the bacterial iron transport systems. This “Trojan horse strategy” leads to increased periplasmic concentrations and avoids porin-mediated resistance mechanisms [1]. Secondly, the ceftazidime and cefepime related sidechains of cefiderocol provides improved hydrolytic stability against various β-lactamases including carbapenemases [2, 3] (Supplementary Figure S1). Indeed, cefiderocol was shown to be hydrolyzed several orders of magnitudes less by various carbapenem hydrolyzing β-lactamases, such as KPC-3, NDM-1 and VIM-2, compared to similar β-lactam drugs such as ceftazidime [2]. Clinical resistance to cefiderocol has mainly been associated with mutations in iron transporter systems [4-6]. However, a two amino acid deletion in the R2 loop of AmpC of a clinical *Enterobacter* spp. isolate led to reduced susceptibility towards cefiderocol as well as ceftazidime-avibactam [7, 8]. Moreover, KPC variants conferring increased ceftazidime-avibactam resistance resulted in cross-resistance against cefiderocol [9]. Further, increased copy number and expression of *bla*_NDM-5_ in *Escherichia coli* was shown to be associated with the development of cefiderocol resistance [10]. Additionally, the synergistic effects between cefiderocol with β-lactamase inhibitors indicate that the expression of β-lactamases might play a crucial role in cefiderocol resistance development [11]. Thus, there is a clear potential for the selection of new or pre-existing β-lactamase variants exhibiting increased resistance towards cefiderocol.

However, a general understanding of the contribution and evolvability of these enzymes to changes in bacterial cefiderocol resistance is currently still lacking. Here, we provide a systematic study addressing this knowledge gap by asking to which extent, the expression of clinical and contemporary β-lactamases from different Ambler classes play a role in the evolution of cefiderocol resistance? Moreover, can the exposure to cefiderocol lead to cross-resistance and re-sensitization (collateral sensitivity) towards other β-lactams and β-lactam-inhibitor combinations? To this end, five β-lactamases, KPC-2 and CTX-M-15 (Ambler class A), NDM-1 (class B), CMY-2 (class C), and OXA-48 (class D) were expressed in *Escherichia coli* using a low-copy number vector system (∼15 copies per cell) since these β-lactamases are often plasmid-associated [12]. First, changes in susceptibility due to the expression of β-lactamases were analyzed by determining the inhibitory concentration 50% (*IC*_50_) and the standard minimum inhibitory concentration (MIC) against a panel of different β-lactams. Next, we used directed evolution to probe the evolutionary potential of these β-lactamases by constructing mutational libraries and selecting clones with increased cefiderocol resistance. We show that β-lactamases of various Ambler classes affect cefiderocol susceptibility and demonstrate evolvability to further cefiderocol resistance, which is often associated with collateral effects.

## MATERIALS AND METHODS

### Antibiotics and other agents

Cefiderocol was purchased from Shionogi (Osaka, Japan). If not otherwise stated, other antibiotics and media were purchased from Sigma-Aldrich (St. Louis, MO, USA). Strains used and constructed in this study are summarized in Supplementary Table 1. Restriction enzymes, DNA polymerases and T4 ligase were purchased from ThermoFisher (Waltham, MA, USA). Primer sequences used in this study are summarized in Supplementary Table 2.

### Strain construction

Previously, we constructed a low-copy number vector (pUN) with a chloramphenicol resistance marker (pA15 origin with ∼10 copies per cell) [13, 14]. The chloramphenicol marker carried a *Nco*I restriction site which was silenced by site directed mutagenesis using GoldenGate cloning. In brief, whole vector amplification was performed with Phusion polymerase and primers P9/P10 (Supplementary Table 2). The PCR product was digested using *Lgu*I and *Dpn*I. Recirculation was performed using a T4 ligase and MP21-05 (*E. coli* E.cloni^®^ 10G) was transformed with the ligated product. Clones were selected on LB plates containing 25 mg/L chloramphenicol and verified by Sanger sequencing (Genewiz, Leipzig, Germany).

This modified vector allowed us to sub-clone all β-lactamases, using a *Nco*I restriction site at the start codon and the *Xho*I restriction site directly after the stop codon. The gene sequences of *bla*_CMY-2_, *bla*_CTX-M-15_, and *bla*_NDM-1_ were synthesized by Genewiz (Leipzig, Germany) according to the gene sequences NG_048935.1, NG_048814.1 and NG_049326.1, respectively. *Bla*_OXA-48_ and *bla*_KPC-2_ originated from *E. coli* 50579417 and *Klebsiella pneumoniae* K47-25, respectively [15, 16].

Primers were designed, replacing the native *Nde*I restriction site at the start codon of the β-lactamase genes with a *Nco*I cutting site by inserting a glycine after the starting methionine amino acid (Supplementary Table 2). For sub-cloning, the vector backbone was amplified using the primers P3/P4 and Phusion polymerase. Similarly, *bla*_OXA-48_ (P1/P2), *bla*_KPC-2_ (P41/42), *bla*_CMY-2_ (P52/53), *bla*_CTX-M-15_ (P48/49) and *bla*_NDM-1_ (P50/51) were amplified, followed by a *Nco*I/*Xho*I digestion. The digested vector backbone and insert were T4 ligated and MP21-05 was transformed with the ligated product. The *Nco*I and *Not*I restriction site within *bla*_KPC-2_ were removed using primers P43F/R and P44F/R, respectively, and GoldenGate cloning as described above.

After selective plating on cefiderocol agar plates, mutant alleles were amplified using primers P7/P8, sub-cloned into an isogenic pUN vector backbone and transformed into MP21-05 to exclude mutational effects outside of the target gene. The corresponding single and double mutants, which could not be sub-cloned directly, were constructed by GoldenGate cloning, as described above, using the primers in Supplementary Table 2. All changes were confirmed by Sanger sequencing.

### Directed evolution and selective plating

Mutational libraries were constructed by error-prone PCR using 10 ng vector DNA, GoTag DNA polymerase (Promega, Madison, WI, USA), 25 mM MgCl_2_ (Promega), 10 μM of primers P7/P8 and either 50 μM oxo-dGTP or 1 μM dPTP. PCR products were DpnI digested for 1 h at 37°C. 5 ng of each product was used for a second PCR, which was performed as described above, but without mutagenic nucleotides. The second PCR product was then digested using *Nco*I and *Xho*I and ligated in a 1:3 ratio with the digested and purified vector backbone. MP21-05 was transformed with the ligation mixture, recovered in LB broth for 1 h at 37°C and plated on 25 mg/L chloramphenicol LB agar plates. Library sizes were determined by cell counts and mutation frequencies were determined using Sanger sequencing (Genewiz, Leipzig, Germany). Sequences were aligned using ESPript (v. 3) [17]. The MP21-05 cultures harboring the corresponding mutational libraries of β-lactamase genes were plated on LB agar plates containing increasing concentrations of cefiderocol and grown over night at 37°C. Colonies grown were recovered on plates with the highest cefiderocol concentration and their genotype characterized by Sanger sequencing (Genewiz, Leipzig, Germany).

### Dose-response curves and MIC determination

Dose response curves were determined and their *IC*_*50*_ values calculated using GraphPad Prism (v. 9) as previously published [14]. MICs were determined by broth microdilution using in-house-designed premade Sensititre microtiter plates (TREK Diagnostic Systems/Thermo Fisher Scientific, East Grindstead, UK) according to the manufacturer’s instruction. The plates were incubated statically for 20 h at 37°C.

## RESULTS

### Evolution of β-lactamase mediated cefiderocol resistance

To comparatively study the effect of different β-lactamases on cefiderocol resistance development, we expressed five β-lactamase genes (*bla*_CMY-2_, *bla*_CTX-M-15_, *bla*_NDM-1_, *bla*_OXA-48_ and *bla*_KPC-2_) in a low-copy number vector in an isogenic *E. coli* E.cloni^®^ 10G (MP21-05) background and determined changes in cefiderocol MICs (Table 1). We found that the expression of OXA-48 and KPC-2 conferred non or marginal (4-fold) decreases in susceptibility towards cefiderocol, respectively. In contrast, CMY-2, CTX-M-15 and NDM-1 substantially decreased cefiderocol susceptibility by 16-, 8- and 32-fold, respectively. Thus, our data show that the expression of contemporary and clinically relevant β-lactamases can be critical and contribute to cefiderocol resistance, which is in-line with previous observations [9, 11, 18].

**Table 1:**
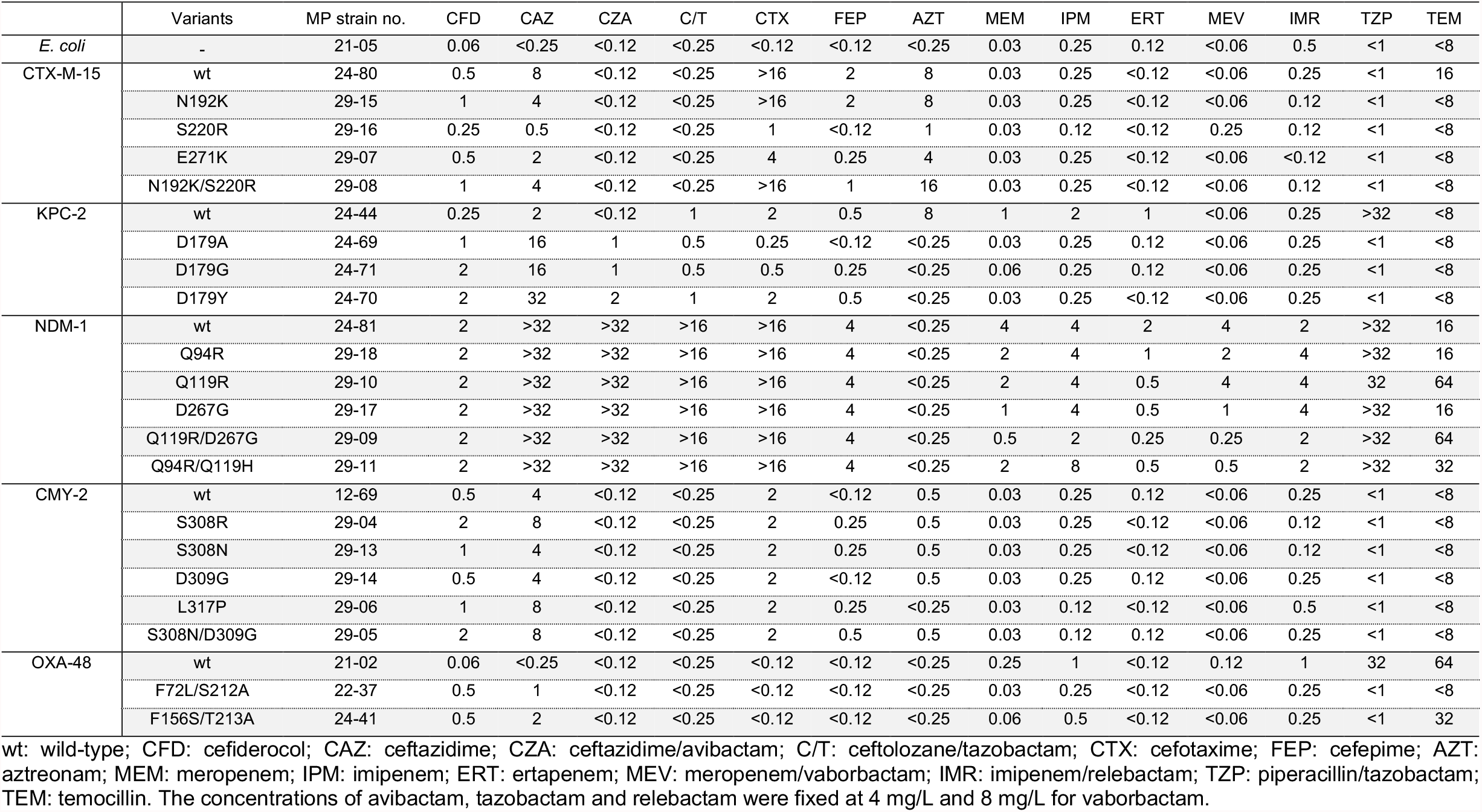
MIC determination. Enzymes are expressed in *E. coli* E.cloni^®^ (MP21-05) and MIC values are reported in mg/L.

Further, the observed effect on cefiderocol susceptibility by the β-lactamases suggests an evolutionary potential for the adaption toward increasing cefiderocol concentrations. To study this, we created mutational libraries, comprising at least 5000 mutants of each β-lactamase, using error prone PCR with an average mutation rate of 1 to 2 mutations per gene. Mutational libraries were selected on agar plates with cefiderocol concentrations 2 to 16-fold above their wild-type MICs. Up to eight colonies were randomly selected per β-lactamase from plates containing the highest cefiderocol concentration, and changes in the target genes were characterized by Sanger sequencing. Among isolated variants, we selected a subset of single and double mutants, with amino acid changes either close to the active site, in structural elements important in substrate specificity (e.g., Ω loop) or described in naturally evolving variants for subsequent characterization (Supplementary Figures 2 to 6): two OXA-48 double mutants (F72L/S212A and F156S/T213A), three KPC-2 single mutants (D179A/G/Y), two single (S308R and L317P) and one CMY-2 double mutant (S308N/D309G), two NDM-1 double mutants (Q119R/D267G and Q94R/Q119H), as well as one single (E271K) and one double CTX-M-15 mutant (N192K/S220R). In addition, to identify the contribution of each individual amino acid change within the selected double mutants, the corresponding single mutants were constructed (Supplementary Table S1).

Standard MIC assays have a limited resolution and may not capture marginal changes in susceptibility that, from an evolutionary perspective, have shown to be crucial for the selection of antibiotic resistance [14, 19]. To provide an increased resolution to our susceptibility measurements, we determined the cefiderocol susceptibility changes using dose response curves (Figure 1) and calculated the corresponding *IC*_50_ values (Table 2). We found that, with the acquisition of only one mutation, all β-lactamases evolved to confer significantly increased resistance (herein defined as decreased susceptibility compared to wild-type allele) against cefiderocol where *IC*_50_ values typically increased by 2 to 8-fold (Brown-Forsythe and Welch ANOVAs for samples with different standard deviations, see Supplementary Table 3). Interestingly, our data also show synergy between different single mutants during the evolution of OXA-48 (F72L/S212A and F156S/T213A), CMY-2 (S308N/D309G) and NDM-1 (Q119R/D267G) where the *IC*_50_ values, conferred by the double mutants, were significantly higher than either single mutant alone (Supplementary Table 3). On the contrary, CTX-M-15:E192K and NDM-1:Q94R did not contribute to cefiderocol resistance development in neither single nor double mutant and are thus likely to be hitch-hikers.

**Table 2:** *IC*_50_ determination. SEM represents the standard error based on at least 3 replicates.

| MP Strain no. | Name | $IC_{50}$<br>(mg/L) | SEM<br>(mg/L) |
| --- | --- | --- | --- |
| MP21-05 | *E. coli* E.cloni® | 0.060 | 0.004 |
| MP24-80 | wtCTX-M-15 | 0.653 | 0.075 |
| MP29-15 | N192K | 0.731 | 0.08 |
| MP29-16 | S220R | 1.122 | 0.083 |
| MP29-07 | E271K | 1.339 | 0.08 |
| MP29-08 | N192K/S220R | 1.16 | 0.142 |
| MP24-44 | wtKPC-2 | 0.102 | 0.004 |
| MP24-69 | D179A | 0.338 | 0.034 |
| MP24-71 | D179G | 0.299 | 0.027 |
| MP24-70 | D179Y | 0.558 | 0.037 |
| MP24-81 | wtNDM-1 | 1.087 | 0.172 |
| MP29-10 | Q94R | 1.126 | 0.312 |
| MP29-17 | Q119R | 4.448 | 0.253 |
| MP29-18 | D267G | 2.489 | 0.323 |
| MP29-09 | Q94R/Q119H | 3.683 | 0.238 |
| MP29-11 | Q119R/D267G | 5.284 | 0.418 |
| MP12-69 | wtCMY-2 | 0.216 | 0.031 |
| MP29-04 | S308R | 1.677 | 0.112 |
| MP29-13 | S308N | 1.466 | 0.108 |
| MP29-14 | D309G | 0.807 | 0.083 |
| MP29-06 | L317P | 1.596 | 0.102 |
| MP29-05 | S308N/D309G | 2.06 | 0.096 |
| MP21-01 | wtOXA-48 | 0.058 | 0.003 |
| MP22-05 | F72L | 0.076 | 0.005 |
| MP22-19 | S212A | 0.066 | 0.005 |
| MP22-06 | F156S | 0.148 | 0.016 |
| MP22-07 | T213A | 0.055 | 0.004 |
| MP22-37 | F72L/S212A | 0.237 | 0.021 |
| MP24-41 | F156S/T213A | 0.381 | 0.033 |

wt: wild-type

**Figure 1:**
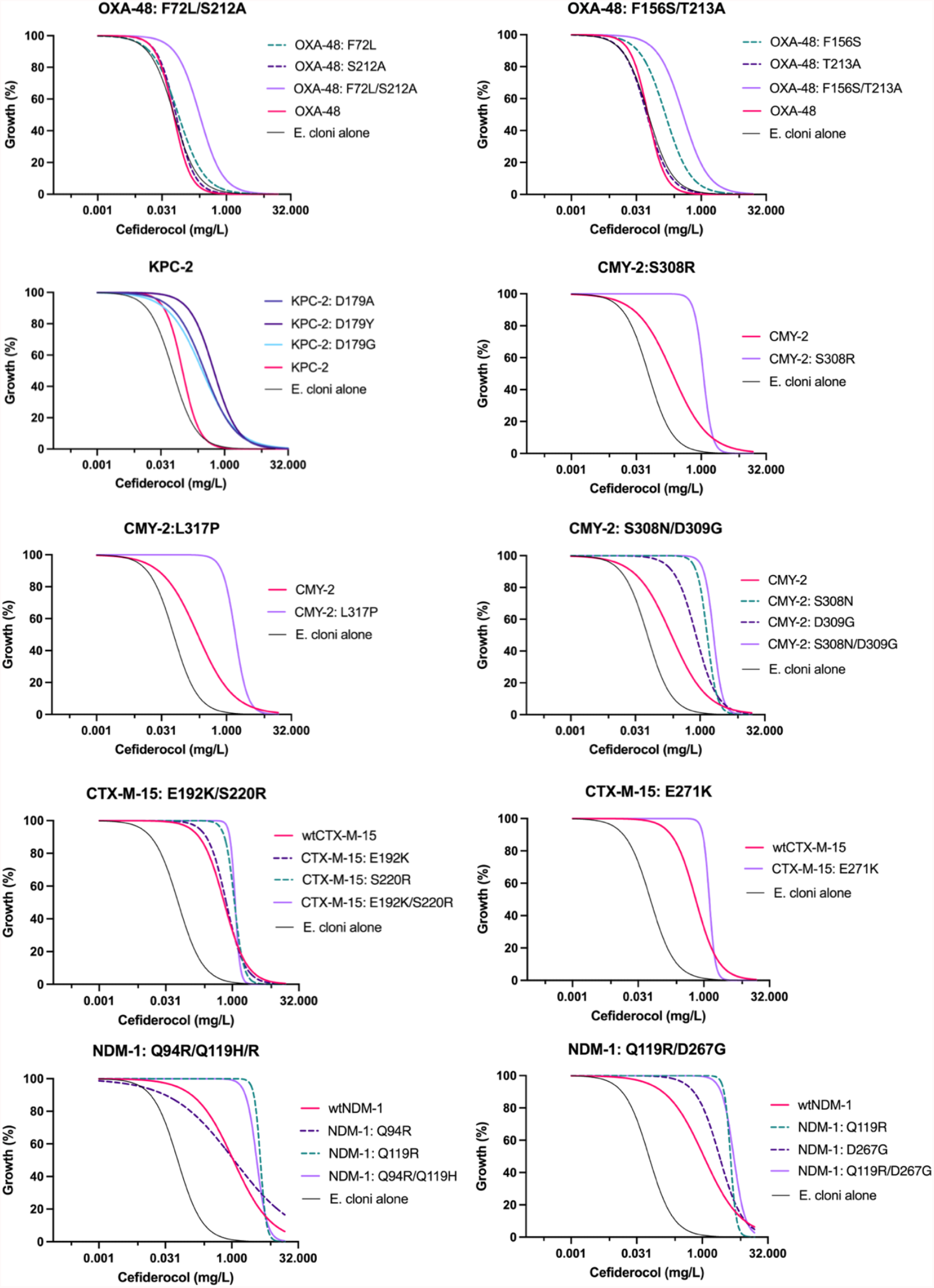
Cefiderocol dose-response curves.

To assess the *IC*_*50*_ changes in a more clinical microbiological context, we further performed standard MIC susceptibility assays. Using this approach, significant cefiderocol MIC differences (>2-fold changes) were only observed for OXA-48, KPC-2 and CMY-2 mutants compared to their wild-type enzymes (Table 1). On the contrary, all mutants of CTX-M-15 and NDM-1 displayed unchanged cefiderocol MIC values, despite their significant changes in *IC*_50_. Taken together, tested β-lactamases of all Ambler classes can evolve to confer decreased susceptibility against cefiderocol, judged by their *IC*_50_ values, while exhibiting cryptic phenotypes from a clinical microbiological point of view (no changes in MIC). However, these marginal changes in resistance have been shown to be highly selectable, especially under low or suboptimal β-lactam concentrations [14] and can provide a gateway for developing clinical resistance [19-22].

### Cefiderocol resistance display changes in collateral susceptibility

Cefiderocol is an oxyimino cephalosporin combining chemical moieties of ceftazidime and cefepime (Supplementary Figure 1). Evolution of β-lactamase mediated resistance towards ceftazidime and ceftazidime combinations with β-lactamase inhibitors, such as avibactam, has been reported to cause collateral changes, e.g., cross-resistance and collateral sensitivity in different enzymes, including KPC and OXA-48 [9, 13]. To understand whether collateral effects occur during the evolution towards cefiderocol resistance, we determined MICs against a panel of different β-lactams, covering all β-lactam classes (Table 1). Our MIC data show that a 4 to 8-fold increase in cefiderocol MIC in OXA-48 and KPC-2 mutants caused the development of strong cross-resistance against ceftazidime, with ceftazidime MICs elevated by 4- to 16-fold. In addition, all three selected KPC-2 mutants exhibited cross-resistance with >8- to >16-fold increased MIC values against ceftazidime-avibactam. These observations are in-line with previous studies where the selection of KPC-2 on ceftazidime-avibactam caused the emergence of KPC-2 mutants conferring cross-resistance against cefiderocol [9]. Similarly, the selection of OXA-48 against ceftazidime resulted in mutants identical or similar (e.g., F72L and F156C/V) to the ones identified in this study [14] suggesting that the exposure to either ceftazidime or cefiderocol causes functional cross-resistance in both KPC-2 and OXA-48. No effect of the OXA-48 mutations on ceftazidime-avibactam resistance development was found and cross-resistance to other cephalosporins, such as cefepime and cefotaxime, was not detected. In addition, no cross-resistance was observed towards ceftazidime-avibactam in CMY-2 and CTX-M-15.

We also observed that evolved cefiderocol resistance comes with a range of significant evolutionary trade-offs. For all three carbapenemases (OXA-48, KPC-2 and NDM-1), we found significant collateral sensitivities towards carbapenems. This was particularly true for the serine carbapenemases OXA-48 and KPC-2, where cefiderocol resistance development caused strong collateral sensitivity effects with reduced carbapenem MICs. The strongest effect was seen for meropenem with a MIC reduction of up to 32-fold in the KPC mutants. A smaller collateral sensitivity effect was observed within the NDM-1:Q119R/D267G mutant where the meropenem MIC was reduced by 8-fold. In addition, in both OXA-48 and KPC-2, mutations resulted in a >32-fold decrease in piperacillin-tazobactam MIC. Other collateral sensitivity changes with MIC reductions >2-fold include ceftazidime (CTX-M-15:S220R and E271K), cefotaxime (KPC-2:D179A/G, CTX-M-15:S220R and CTX-M-15:E271K) and aztreonam (CMY-2:L317P, KPC-2 D179x and CTX-M-15:S220R).

## DISCUSSION

There have been observations that β-lactamases may impact the susceptibility and evolution of cefiderocol resistance [7-11, 18]. Indeed, we showed that the expression of wild-type β-lactamases from various Ambler classes can significantly alter cefiderocol susceptibility in *E. coli* (Table 1). Beyond that, we probed the evolutionary potential of all tested β-lactamases to adapt toward increasing cefiderocol resistance (Table 2). With the acquisition of only one to two mutations, all β-lactamases evolved to confer increased resistance against cefiderocol. Interestingly, we observed that the extent by which cefiderocol resistance developed was highly dependent on the initial wild-type β-lactamase activity. Enzymes with an initial low-level resistance profile against cefiderocol, such as OXA-48 and KPC-2, showed the highest improvement. On the contrary, enzymes with an initial higher level of activity against cefiderocol, such as CMY-2, NDM-1, and CTX-M-15, demonstrated substantial less improvements (Table 1 and 2). Indeed, selected mutants for these enzymes did not significantly improve cefiderocol resistance judged by a standard MIC assay. However, we found that most of the mutants were able to significantly improve resistance measured in an *IC*_50_ set-up. Such cryptic changes in susceptibility have been previously described to play an important role in the evolution of β-lactamases and are highly selectable, especially under sub-optimal β-lactam concentrations [14, 19]. We acknowledge the fact that only single and double mutants were studied, and further work needs to be done to explore the full evolutionary potential of these enzymes.

Evolution of β-lactamase mediated resistance to ceftazidime-avibactam and cefepime has been shown to concurrently cause cross-resistance or reduced susceptibility to cefiderocol [7-9]. Here, we observed a similar phenomenon where cefiderocol and ceftazidime resistance increased parallelly to the same extent within OXA-48 and KPC-2 (Table 1) and ceftazidime-avibactam resistance in KPC-2. No cross-resistance against other oxyimino cephalosporins or β-lactams was identified indicating that the structural similarity between cefiderocol and ceftazidime plays an important role for the development of cross-resistance.

In contrast, widespread collateral sensitivity against other β-lactams, including carbapenems and penicillin-inhibitor combinations, was found in mutants with increased cefiderocol resistance (Table 1). We observed the strongest trade-offs during cefiderocol resistance development of OXA-48 and KPC-2 against carbapenems and aztreonam (KPC-2). Such collateral sensitivity/functional trade-offs can open the path for alternative treatment strategies, and they have been successfully exploited in the clinical setting with a carbapenem-β-lactamase inhibitor combination against ceftazidime-avibactam and cefiderocol resistant *K. pneumoniae* harboring the natural KPC-31[23]. However, the molecular causes of these collateral effects remain poorly understood. A study on the ceftazidimase OXA-163, which possesses lower carbapenem activity compared to OXA-48, suggests that molecular evolution shapes drug incompatibility, resulting in multiple binding modes that give rise to these trade-offs.

F72 and F156 in OXA-48 have been previously characterized as mutational hotspot allowing for marginally increased ceftazidime hydrolysis [13]. Here, we re-identified mutations at these positions (F72L and F156S) showing their involvement in cefiderocol resistance development. While OXA-48:F72L was reported in environmental samples [24, 25], most characterized OXA-48-like variants, which confer increased ceftazidime resistance, exhibit multiple amino acid deletions within the ß5-ß6 loop [26]. It remains to be determined whether these variants, such as OXA-163, also confer increased resistance against cefiderocol. In contrast, the D179x mutations within the Ω-loop of KPC-type have been described in naturally evolving enzymes (KPC-78, KPC-86 and KPC-31; Supplementary Figure 3). For CMY-2, amino acid changes clustered round the R2 loop which has been shown to be host to the R2-side chain of β-lactam drugs [27]. Consequently, mutations and deletions within the R2 loop have been associated with increased resistance towards cephalosporins, such as cefepime and ceftazidime [28, 29]. Also here, several of the mutations/positions reported in this study have been associated with naturally evolving variants (e.g., CMY-133 and CMY-17). This underlines the fact that variants conferring improved cefiderocol resistance are already present in clinical isolates. In addition, these enzymes are encoded on transferable plasmids allowing these genes to spread, a process that may be facilitated by the increasing usage of cefiderocol.

Taken together, this study provides a proof-of-principle showing that the expression of β-lactamases from various Ambler classes can substantially contribute to cefiderocol resistance and that many β-lactamases possess the evolutionary potential to adapt to increasing cefiderocol concentrations under laboratory conditions. Similar to other cephalosporins, this evolutionary process comes with collateral effects against β-lactam drugs including both cross-resistance and re-sensitization.

## ACKNOLEGEMENTS

We are very grateful to Hanna-Kirsti Schrøder Leiros, Department of Chemistry, UiT The Arctic University of Norway for providing the cefiderocol used in this study.

