## Supplementary information for "Evolution of β-lactamase mediated cefiderocol resistance"

### Supplementary Table 1: Strains used and constructed in this study

| Strain no.* | Comment | Reference |
|---|---|---|
| K47-25 | Host strain of *bla*$_{KPC-2/\textit{NcoI/NotI}}$ | [1] |
| 50579417 | Host strain of p50579417_3_OXA-48 | [2] |
| MP13-01/ MP08-01 | K12 MG1655 | Uppsala/Lisbon University |
| MP21-05 | *E. coli* E.cloni® 10G | Lucigen |
| MP13-06 | MP13-01 transformed with p50579417_3_OXA-48 | [3] |
| MP08-61 | MP08-01 carrying pUN-*bla*$_{OXA-48}$ | [4] |
| MP21-01 | MP21-05 carrying pUN-*bla*$_{OXA-48}$ | This study |
| MP12-70 | MP21-05 carrying pUN-*bla*$_{KPC-2/\textit{NcoI/NotI}}$ | This study |
| MP24-43 | MP21-05 carrying pUN-*bla*$_{KPC-2/\textit{NcoI}}$ | This study |
| MP24-44 | MP21-05 carrying pUN-*bla*$_{KPC-2}$ | This study |
| MP24-80 | MP21-05 carrying pUN-*bla*$_{CTX-M-15}$ | This study |
| MP12-69 | MP21-05 carrying pUN-*bla*$_{CMY-2}$ | This study |
| MP24-81 | MP21-05 carrying pUN-*bla*$_{NMD-1}$ | This study |
| MP21-02 | MP21-05 carrying mutational library of pUN-*bla*$_{OXA-48}$ | This study |
| MP24-45 | MP21-05 carrying mutational library of pUN-*bla*$_{KPC-2}$ | This study |
| MP29-01 | MP21-05 carrying mutational library of pUN-*bla*$_{CTX-M-15}$ | This study |
| MP29-03 | MP21-05 carrying mutational library of pUN-*bla*$_{CMY-2}$ | This study |
| MP29-02 | MP21-05 carrying mutational library of pUN-*bla*$_{NMD-1}$ | This study |
| MP22-05 | MP21-05 carrying pUN-*bla*$_{OXA-48: F72L}$ | This study |
| MP22-19 | MP21-05 carrying pUN-*bla*$_{OXA-48: F156S}$ | This study |
| MP22-06 | MP21-05 carrying pUN-*bla*$_{OXA-48: S212A}$ | This study |
| MP22-07 | MP21-05 carrying pUN-*bla*$_{OXA-48: T213A}$ | This study |
| MP22-37 | MP21-05 carrying pUN-*bla*$_{OXA-48: F72L/S212A}$ | This study |
| MP24-41 | MP21-05 carrying pUN-*bla*$_{OXA-48\ F156S/T213A}$ | This study |
| MP24-69 | MP21-05 carrying pUN-*bla*$_{KPC-2: D179A}$ | This study |
| MP24-71 | MP21-05 carrying pUN-*bla*$_{KPC-2: D179G}$ | This study |
| MP24-70 | MP21-05 carrying pUN-*bla*$_{KPC-2: D179Y}$ | This study |
| MP29-15 | MP21-05 carrying pUN-*bla*$_{CTX-M-15: N192K}$ | This study |
| MP29-16 | MP21-05 carrying pUN-*bla*$_{CTX-M-15: S220R}$ | This study |
| MP29-07 | MP21-05 carrying pUN-*bla*$_{CTX-M-15: E271K}$ | This study |
| MP29-08 | MP21-05 carrying pUN-*bla*$_{CTX-M-15: N192K/S220R}$ | This study |
| MP29-04 | MP21-05 carrying pUN-*bla*$_{CMY-2: S308R}$ | This study |
| MP29-13 | MP21-05 carrying pUN-*bla*$_{CMY-2: S308N}$ | This study |
| MP29-14 | MP21-05 carrying pUN-*bla*$_{CMY-2: D309G}$ | This study |
| MP29-06 | MP21-05 carrying pUN-*bla*$_{CMY-2: L317P}$ | This study |
| MP29-05 | MP21-05 carrying pUN-*bla*$_{CMY-2: S308N/D309G}$ | This study |
| MP29-18 | MP21-05 carrying pUN-*bla*$_{NMD-1: Q94R}$ | This study |

| | | |
|---|---|---|
| MP29-10 | MP21-05 carrying pUN-*bla*$_{NMD-1:\ Q119R}$ | This study |
| MP29-17 | MP21-05 carrying pUN-*bla*$_{NMD-1:\ D267G}$ | This study |
| MP29-11 | MP21-05 carrying pUN-*bla*$_{NMD-1:\ Q94R/Q119H}$ | This study |
| MP29-09 | MP21-05 carrying pUN-*bla*$_{NMD-1:\ Q119R/D267G}$ | This study |

*All strains are *E. coli* species except of K47-25 which is *Klebsiella pneumoniae.*

### Supplementary Table 2: Primers used in this study

| No. | Name | 5´- 3´ | Reference |
|---|---|---|---|
| P1 | *Nco*I-OXA-48 | GCTTCCATGGGACGTGTATTAGCCTTATCGGC | This study |
| P2 | *Xho*I-OXA-48 | GCTTCTCGAGCTAGGGAATAATTTTTTCCTGTTTGAGC | This study |
| P3 | *Nco*I-pUN | GCTTTCCCATGGATGTTTTTCCTCCTTATGTTAAGCTTACTCAG | This study |
| P4 | *Xho*I-pUN | GCTTCATATGTTTTTCCTCCTTATGTTAAGCTTACTCAG | This study |
| P9 | Silencing-*Nco*I-CMY-2-F | TTTTTTGCTCTTCTGCCCATCGTGAAAACGGGGG | This study |
| P10 | Silencing-*Nco*I-CMY-2-R | TTTTTTGCTCTTCGGGCAAATATTATACGCAAGGCG | This study |
| P15F | OXA-48-F72L | TTTTTT GCTCTTC G CAT C TACC CTG AAAATTCCCAATAGCTT | This study |
| P15R | OXA-48-F72L | TTTTTT GCTCTTC GATGC GGGTAAAAATGCTTG | This study |
| P16F | OXA-48-S212A | TTTTTTGCTCTTCACTGGATACGCGACTAGAATCGAACCTAAGATTGG | This study |
| P16R | OXA-48-S212A | TTTTTTGCTCTTC CCAGTTTTAGCCCGAATAATATAGTCACC | This study |
| P28R | OXA-48-F156S | TTTTTT GCTCTTC TCTACATTGCCCGAAATGTC | This study |
| P41 | *Nco*I-KPC-2 | TTTTTTCCATGGGATCACTGTATCGCCGTCTAGT | This study |
| P42 | *Xho*I-KPC-2 | TTTTTTCTCGAGTTACTGCCCGTTGACGCCC | This study |
| P43F | Silencing-*Nco*I-KPC-2 | TTTTTTGCTCTTCAGACTCGCGCTTGAGGGATTGGGCG | This study |
| P43R | | TTTTTTGCTCTTCAGTCTAGCCGCAGCGGCGAT | This study |
| P44F | Silencing-*Not*I-KPC-2-F | TTTTTTGCTCTTCGAGCTGTCCGCCGCCGCCGTGCAATACAG | This study |
| P44R | | TTTTTTGCTCTTCAGCTCCGCCACCGTCATGCC | This study |
| P45F | OXA-48-F156S | TTTTTT GCTCTTC GTAGA CAGT AGC TGGCTCGACGGTGGT | This study |
| P48 | *Nco*I-CTX-M-15 | GCTTCCATG GGAGTTAAAAAATCACTGCGCC | This study |
| P49 | *Xho*I-CTX-M-15 | GCTTCTCGAGTTACAAACCGTCGGTGACG | This study |
| P50 | *Nco*I-NDM-1 | GCTTCCATGGGAGAATTGCCCAATATTATGCACCC | This study |
| P51 | *Xho*I-NDM-1 | GCTTCTCGAGTCAGCGCAGCTTGTCGGCC | This study |
| P52 | *Nco*I-CMY-2 | GCTTCCATGGGAATGAAAAAATCGTTATGCTGCG | This study |
| P53 | *Xho*I-CMY-2 | GCTTCTCGAGTTATTGCAGCTTTTCAAGAATGCGCC | This study |
| P59F | CMY-2-S308N | TTTTTTGCTCTTCATCAACGGCAACGACAGCAAAGTGG | This study |
| P60F | CMY-2-D309G | TTTTTTGCTCTTCATCAACGGCAGCGGCAGCAAAGTGG | This study |
| P61R | CMY-SN/DG | TTTTTTGCTCTTCTTGATGATCGAATCAGCTTTCAGCGG | This study |
| P62F | CTX-M-15-N192K | TTTTTTGCTCTTCCTCTGCGGAAGCTGACGCTGGGTAAAG | This study |
| P62R | | TTTTTTGCTCTTCCAGAGTTTGCGCCATTGCCC | This study |
| P63F | CTX-M-15-S220R | TTTTTTGCTCTTCCGGTGCAGCGCGCATTCAGGCTGGACTGC | This study |
| P63R | | TTTTTTGCTCTTCCACCGGTGGTATTGCCTTTCAT | This study |
| P64F | NDM-1-D267G | TTTTTTGCTCTTCTCCGCCCCCGGTAGCCGCGCCGCAATC | This study |
| P64R | | TTTTTTGCTCTTCGCGGAATGGCTCATCACG | This study |
| P65F | NDM1-Q94R | TTTTTTGCTCTTCCAGACCGCCCGGATCCTCAACTGGATC | This study |
| P65R | | TTTTTTGCTCTTCGTCTGGTCATCGGTCCAGG | This study |

### Supplementary Table 3: Brown-Forsythe and Welch ANOVAs

| ANOVA comparisons | 95% CI of difference | Significance | Multiplicity adjusted P values |
|---|---|---|---|
| wtCTX-M-15 *versus* | | | |
| CTX-M-15: E192K | -0.2727 to 0.1170 | ns | 0.6098 |
| CTX-M-15: S220R | -0.6662 to -0.2705 | *** | 0.0007 |
| CTX-M-15: E271K | -0.8804 to -0.4911 | **** | <0.0001 |
| CTX-M-15: N192K/S220R | -0.8117 to -0.2020 | ** | 0.0061 |
| CTX-M-15: N192K/S220R *versus* | | | |
| CTX-M-15: E192K | 0.2157 to 0.292 | ns | 0.8736 |
| CTX-M-15: S220R | 0.1765 to 0.6815 | ** | 0.0062 |
| wtKPC-2 *versus* | | | |
| KPC-2: D179A | -0.3862 to -0.08583 | * | 0.0144 |
| KPC-2: D179G | -0.3163 to -0.07730 | * | 0.0126 |
| KPC-2: D179Y | -0.6222 to -0.2912 | ** | 0.0028 |
| wtNDM-1 *versus* | | | |
| NDM-1: Q94R | -0.7382 to 0.6606 | ns | >0.9999 |
| NDM-1: Q119R | -3.961 to -2.760 | **** | <0.0001 |
| NDM-1: D267G | -2.121 to -0.6815 | ** | 0.0029 |
| NDM-1: Q94R/Q119H | -3.173 to -2.019 | **** | <0.0001 |
| NDM-1: Q119R/D267G | -5.183 to -3.210 | *** | 0.0002 |
| NDM-1: Q94R/Q119H *versus* | | | |
| NDM-1: Q94R | -1.271 to -0.2584 | ** | 0.0087 |
| NDM-1: Q119R | 1.985 to 3.129 | **** | <0.0001 |
| NDM-1: Q119R/D267G *versus* | | | |
| NDM-1: Q119R | 0.08156 to 1.590 | * | 0.0347 |
| NDM-1: D267G | 2.025 to 3.565 | **** | <0.0001 |
| wtCMY-2 *versus* | | | |
| CMY-2: S308R | -1.902 to -1.019 | *** | 0.0002 |
| CMY-2: S308N | -1.676 to -0.8238 | *** | 0.0004 |
| CMY-2: D309G | -0.9272 to -0.2547 | ** | 0.0047 |
| CMY-2: L317P | -1.784 to -0.9762 | *** | 0.0002 |
| CMY-2: S308N/D309G | -2.227 to -1.461 | **** | <0.0001 |
| CMY-2: S308N/D309G *versus* | | | |
| CMY-2: S308N | 0.2012 to 0.9868 | ** | 0.0066 |
| CMY-2: D309G | 0.9081 to 1.598 | **** | <0.0001 |
| wtOXA-48 *versus* | | | |
| OXA-48: F72L | -0.04882 to 0.01351 | ns | 0.2041 |
| OXA-48: F156S | -0.1746 to -0.006220 | * | 0.0412 |
| OXA-48: S212A | -0.03018 to 0.01338 | ns | 0.6103 |
| OXA-48: T213A | -0.01639 to 0.02250 | ns | 0.9829 |
| OXA-48: F72L/S212A | -0.2905 to -0.06823 | * | 0.0131 |
| OXA-48: F156S/T213A | -0.4954 to -0.1499 | ** | 0.0086 |

| | | | |
|---|---|---|---|
| OXA-48: F72L/S212A *versus* | | | |
| OXA-48: F72L | 0.07627 to 0.2472 | ** | 0.0087 |
| OXA-48: S212A | 0.08596 to 0.2560 | ** | 0.0073 |
| OXA-48: F156S/T213A *versus* | | | |
| OXA-48: F156S | 0.1090 to 0.3555 | ** | 0.0057 |
| OXA-48: T213A | 0.1948 to 0.4567 | ** | 0.0039 |

CI: confidence interval; ns: not significant ($P > 0.05$); * $P \le 0.05$; ** $P \le 0.01$; *** $P \le 0.001$; **** $P \le 0.0001$

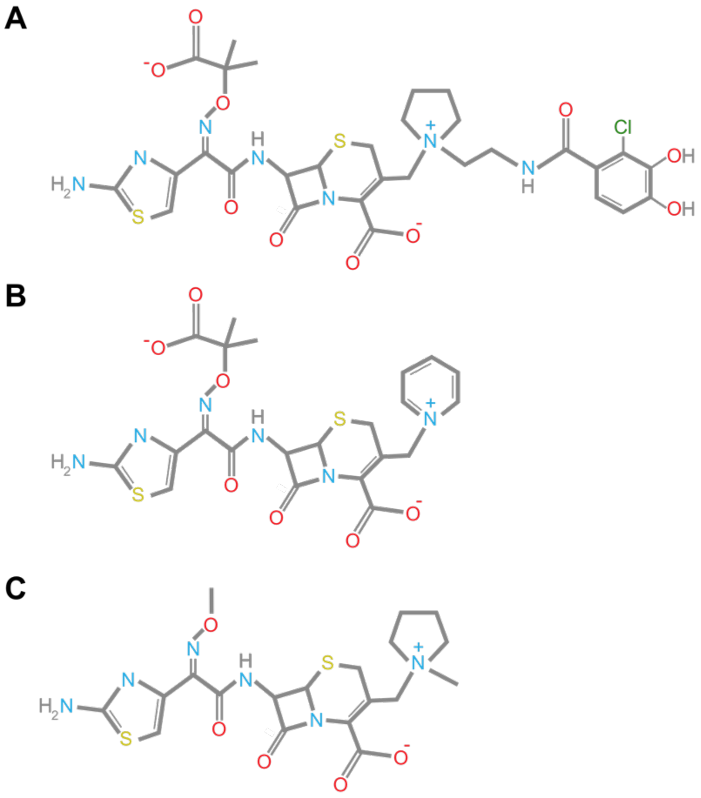


Supplementary Figure 1: Chemical structures of cefiderocol (A), ceftazidime (B) and cefepime (C).

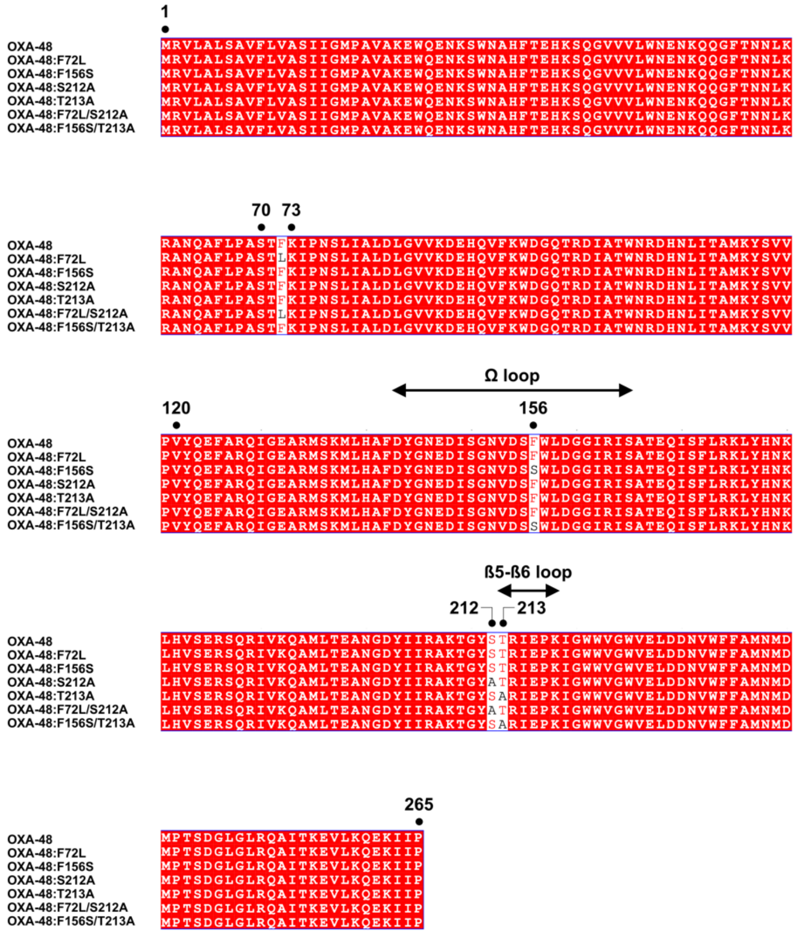

Supplementary Figure 2: OXA-48 sequence alignment with numbering compared to OXA-48, including the active site serine, lysine, and valine at positions 70,73 and 120, respectively. Mutations were found at positions 72,156, 212 and 213. The location of the Ω and ß5-ß6 loops is shown with the arrows.

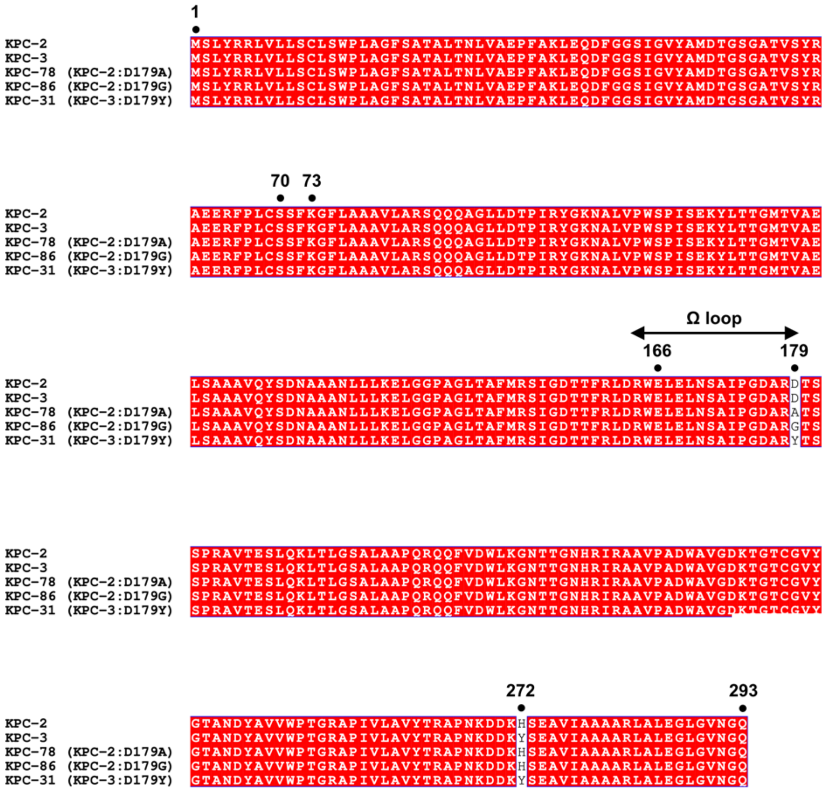


Supplementary Figure 3: KPC-2 alignment sequence alignment according to the class A numbering system [5] including the active site serine, lysine and glutamate at positions 70,73 and 166, respectively. Mutations were found at position 179 within the Ω loop.

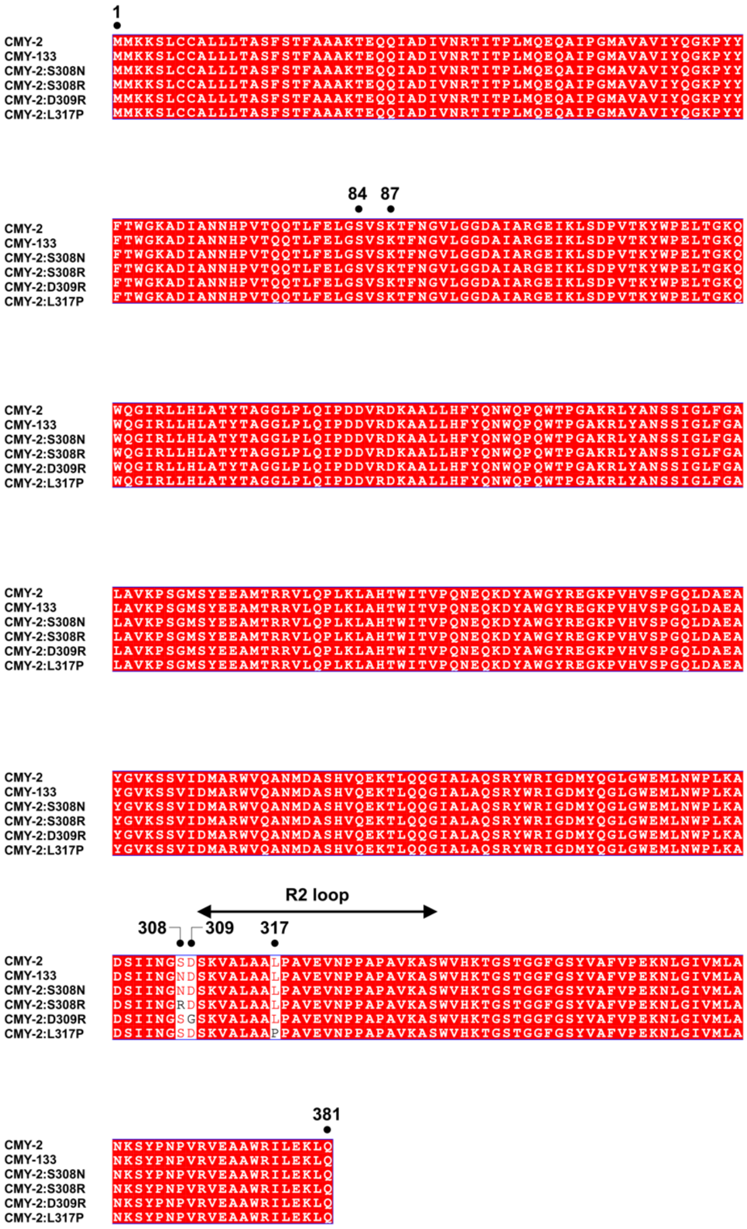

Supplementary Figure 4: CMY-2 sequence alignment compared to CMY-2, including the active site serine and lysine at positions 84 and 87, respectively. Mutations were found at positions either within or close to the R2 loop.

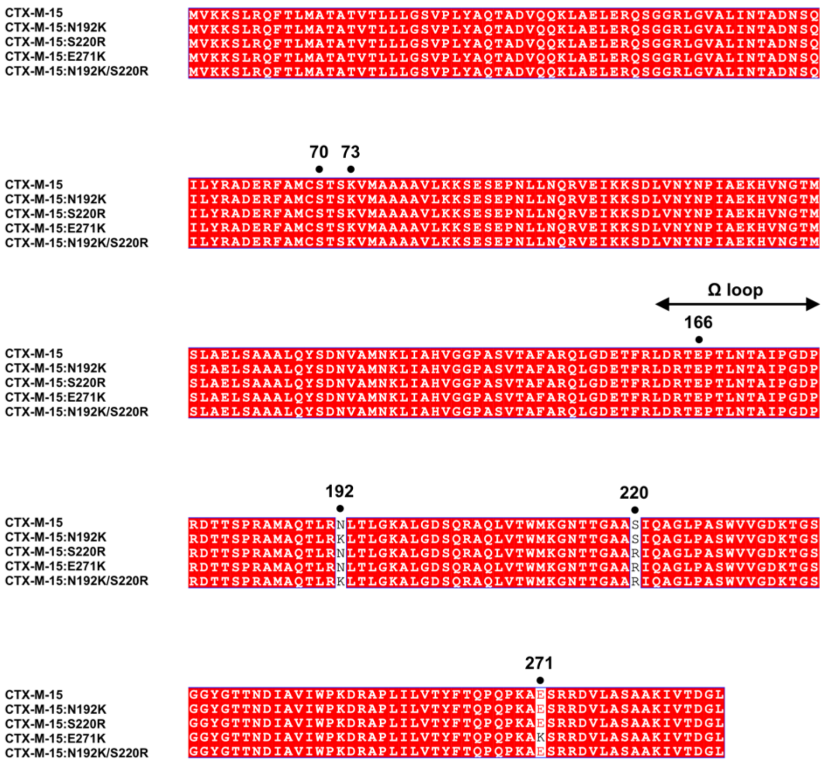


Supplementary Figure 5: CTX-M-15 sequence alignment according to the class A numbering system [5] including the active site serine, lysine and glutamate at positions 70, 73 and 166, respectively. Mutations were found at positions 192, 220 and 271.The position of the Ω loop is indicated with an arrow.

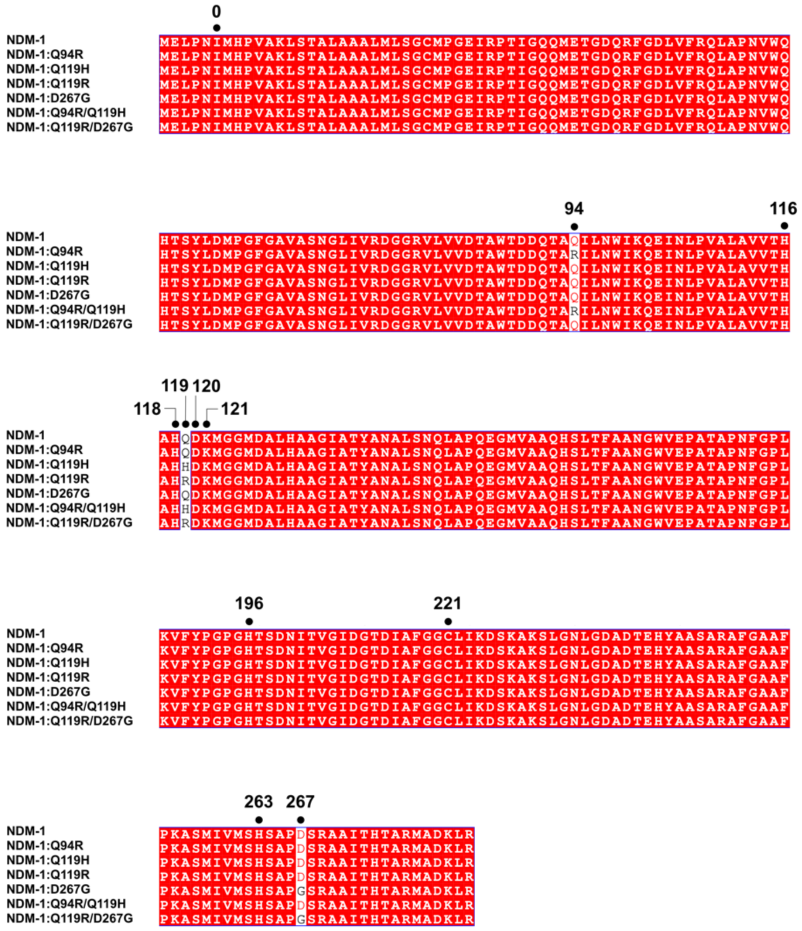


Supplementary Figure 6: NDM-1 sequence alignment according to the metallo-β-lactamase numbering system [6] demonstrating the discovered mutation at position 94,119 and 267. Positions of the metal binding site 1 (116, 118 and 196) and 2 (120,121 and 263) are further numbered.
